# Rosetta Custom Score Functions Accurately Predict ΔΔG of Mutations at Protein-Protein Interfaces Using Machine Learning

**DOI:** 10.1101/2020.03.17.996066

**Authors:** Sumant Shringari, Sam Giannakoulias, John J. Ferrie, E. James Petersson

## Abstract

Protein-protein interfaces play essential roles in a variety of biological processes and many therapeutic molecules are targeted at these interfaces. However, accurate predictions of the effects of interfacial mutations to identify “hotspots” have remained elusive despite the myriad of modeling and machine learning methods tested. Here, for the first time, we demonstrate that nonlinear reweighting of energy terms from Rosetta, through the use of machine learning, exhibits improved predictability of ΔΔG values associated with interfacial mutations.

---

Protein-protein interactions mediate many essential biological processes. Cellular signaling, spatial and temporal regulation, and metabolism are deeply rooted in the formation of higher order protein quaternary structures.^1^ Complex formation is governed by the complementary structural and chemical features displayed by residues at the protein-protein interface, and mutations of these residues are highly correlated with dysfunction and disease.^2^ Moreover, the development of new biomaterials and catalysis strategies are largely dependent on the binding affinity of the involved protein partners.^3, 4^ Therefore, a computational model capable of rapidly and accurately predicting the energy differences (ΔΔGs) associated with mutations would aid in the identification of protein-protein hotspots, providing insight in disease and design.^5, 6^

To date, several approaches have been developed towards the accurate prediction of protein ΔΔG values. These include the use of statistical and contact potentials^7-10^, design of novel sampling schemes,^11, 12^ generation of weighted energy or score functions,^13-16^ and employment of supervised machine learning techniques.^17-21^ Additionally, within the Rosetta Modelling Suite, new sampling schemes, designed to mimic protein motions observed in solution, have afforded increased predictive accuracy.^11, 22^ Though these methods have shown some notable success, there is still a need for a single, generalizable, and facile approach capable of accurately predicting ΔΔG’s of mutations at protein/protein interfaces.

To this end, we envisioned that reweighting of energy terms from Rosetta through machine learning will provide a platform with improved ΔΔG prediction accuracy. The full-atom score function in Rosetta has been repeatedly improved through the introduction of new energy terms and optimization of term weighting. Although Rosetta-based simulations can generate accurate structural models, correlations between the canonical score functions and experimental data remain relatively poor^23^. This suggests that that while the underlying set of terms may produce models with small RMSDs relative to experimental structures, energetically, they require differential weightings for specific applications like ΔΔG prediction. Therefore, we designed the first reported Custom Score Function (CSF, named SRS2020), which is a score function devised purely through the reweighting of Rosetta energy terms for optimal prediction of an experimentally measurable variable of interest. This method allows for the traditional Rosetta score function to be used for structural refinement, while SRS2020 can be used to more accurately predict ΔΔGs. This notion is not entirely novel as protein and small molecule design strategies in Rosetta have used supplementary criteria in the form of classifiers and filters to perform selection based on criteria not encompassed within scoring^24^. However, the approach presented herein is simpler in that it requires no additional terms to be constructed.

To test the utility of this approach, we focused on simulating the SKEMPI 2.0 database (https://life.bsc.es/pid/skempi2/) which is the largest curated database of protein-protein interfacial mutants. This database includes 348 different protein-protein complexes, and ΔΔG values for 6193 unique interfacial mutations of this protein set.^25, 26^ After removing complexes where the mutation location could not be accurately assigned due to ambiguities between the number of protein subunits and were left with 5366 unique mutant complexes. Fig. 1 illustrates the generalized version of our computational protocol employed within PyRosetta.^27^ First, wild-type complexes are cleaned (removal of solvent, ligands, or ions), renumbered, and subjected to initial minimization. Subsequently, mutations are introduced, and both the wild-type and mutant complexes are subjected to another round of structural optimization followed by the computation of Rosetta energies which in turn are fed into a variety of machine learning protocols. Sampling was varied both in the initial, structural stage where only wild-type complexes were considered, as well as in the mutational stage where both the wild-type and mutants were sampled under a uniform scheme. In the structural sampling stage, we assessed the impact of relaxing the input structure compared to using the structure directly from the SKEMPI 2.0 server. Additionally, we tested the impact of local and global sampling during the mutational sampling stage, where either only the mutant residue was packed, or repacking was performed on the entire complex. Lastly, we computed energies for each sampling combination using both REF2015,^28^ the most recently published score function, and BETA_NOV16,^29^ the newest score function available in Rosetta.

**Fig. 1.**
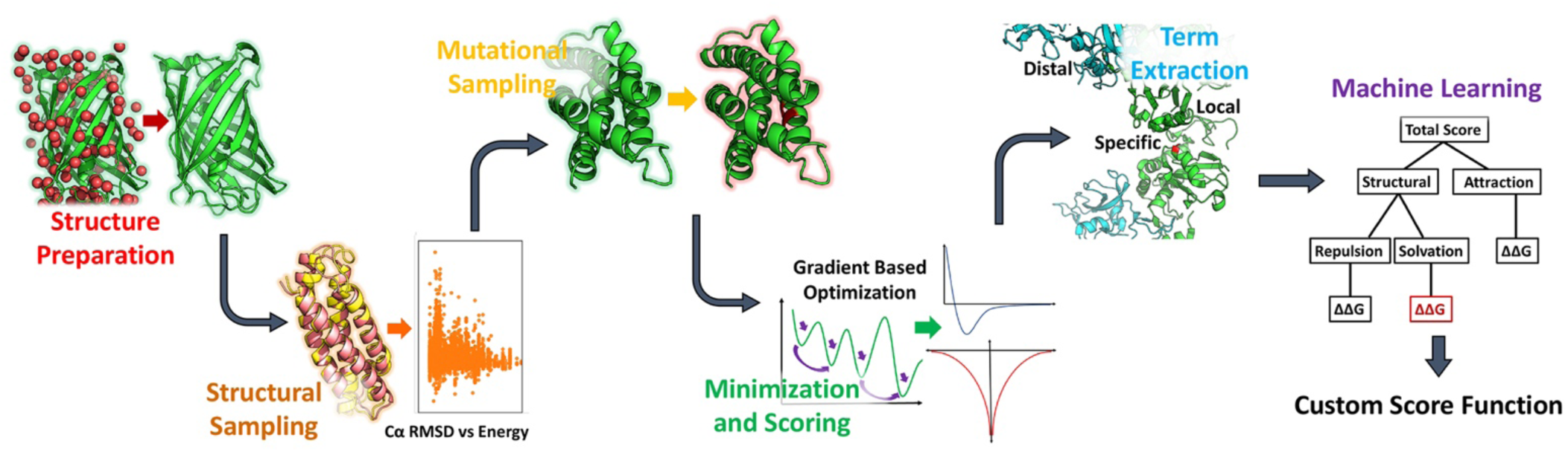
Schematic of computational workflow for developing a custom score function to predict ΔΔG values of mutations at protein/protein interfaces.

We first focused on assessing the performance of the traditional Rosetta score function in predicting mutational ΔΔGs from the SKEMPI 2.0 database. As expected, differences in sampling impacted the correlation of Rosetta total energy scores with ΔΔG. Initial minimization of wild-type complexes, prior to mutational sampling, was found to improve correlation between total Rosetta Energy Units (REUs), and experimental ΔΔG values for both the REF2015 and BETA_NOV16 score functions. This was unsurprising as it is widely recognized that structures determined from crystallographic data require initial relaxation prior to sampling within Rosetta to produce more correlative simulations.^30^ Although we observe an improvement in the predictive capacity following minimization during initial structural sampling, which is likely due to the approximate five REU reduction in the average residue score, we observe only a minimal change in conformation (Supplementary Information, SI, Fig. S1).

Across all sampling and scoring schemes, we see a maximum average RMSD of 0.45 Å compared to input structures from SKEMPI 2.0. Additionally, score values derived from local packing only at the mutation site prior to minimization showed a higher predictive capacity than global repacking following mutation. A Wilcoxon t-test was performed to identify Rosetta energy terms that differed between these simulations. The difference in predictability between these models is likely due to the drastic differences in Lennard-Jones, Dunbrack, and solvation terms produced by these simulations (see SI). Lastly, it is notable that the most recent score function, BETA_NOV16, afforded a higher correlation with experimental data than the benchmarked REF2015 score function. This may be due to increased accuracy in weightings or the additional terms added in the BETA_NOV16 score function.

We sought to improve the predictive power of the models generated by both the REF2015 and BETA_NOV16 score functions using machine learning. Given the results of the structure preparation comparisons, we elected to focus exclusively on improving the predictability of simulations where structures are initially relaxed and only locally sampled following mutation. Multiple linear regression (MLR) is a basic machine learning technique that uses several explanatory variables to predict a single output. Here, those variables are the Rosetta energy terms generated during simulation which will be reweighted to optimize the correlation to experimental ΔΔG. As illustrated in Fig. 2, even this simple MLR approach (Fig. 2B) results in an improved correlation with experimental data compared to the traditional score functions (Fig. 2A). Using both five and ten fold cross validation, we determined that using an MLR improved testing set correlation by a factor of at least 1.58 (SI, Table S1, S2). We also observed, again, that application of the BETA_NOV16 score function displayed a higher predictability than the REF2015 score function. Interestingly, the most important terms in the MLR scoring correlated well with the terms that differed between sampling schemes (SI, Table S3). While this improvement was encouraging, the usability of this reweighted score function is still poor as the Pearson correlation coefficient was only 0.51 and the mean absolute error (MAE) was 1.28 kcal/mol.

**Fig. 2.**
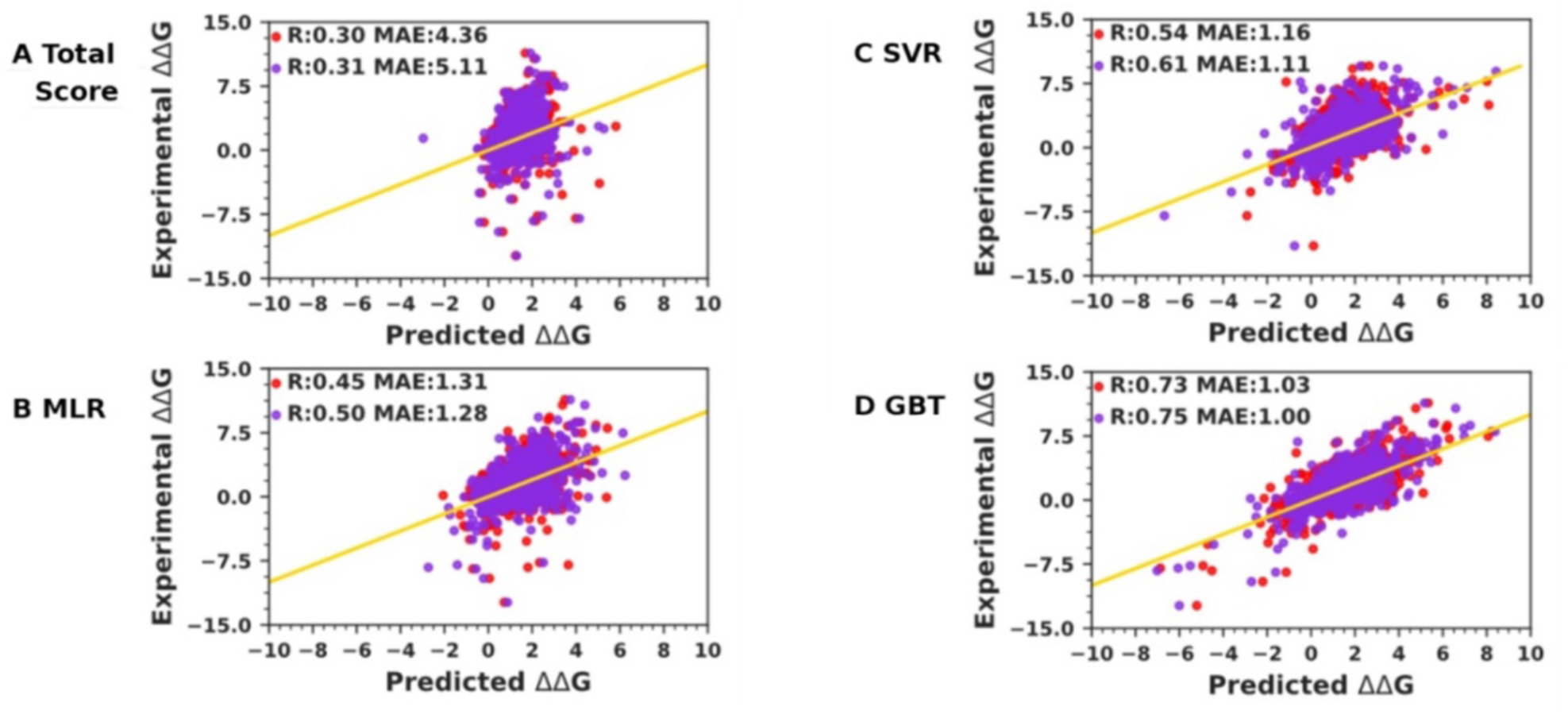
Models for predicting interfacial ΔΔG, REF2015 (red), BETA_NOV16 (purple). Rosetta total score in Rosetta energy units, or REU (A), multiple linear regression (MLR) of Rosetta energy terms (B), polynomial support vector regression (SVR, C), and gradient boosted random forest (GBT) regression (D). MAE: Mean absolute error in kcal/mol.

In order to further improve our ΔΔG predictions, additional inputs were considered, as well as the introduction of more complex machine learning algorithms. Inputs were extracted from simulated structures as the change in energy following sampling. Several different values were computed to capture global vs. specific and local vs. distal differences. Global terms correspond to the change of the total values of the decomposed Rosetta energy terms across the whole mutant and wild-type complexes. Specific values refer directly to the differences between the decomposed Rosetta energy terms of only the mutated and wild-type residues. To distinguish local and distal terms, an 8 Å contacting shell was created around the mutation site. This was used for the calculation of local and distal terms, which correspond to the change in total decomposed scores within or beyond this sphere. In addition to alternative inputs, we employed more sophisticated machine learning algorithms: Kernel Ridge Linear Regression (KRR, see S10, S11), support vector regressions (SVRs), and Gradient Boosted Random Forrest Regression (GBT). Our reasons for choosing these methods are outlined in SI. Training and testing sets were specifically designed to ensure that no mutational redundancy existed between the sets. Curation of training and testing sets in this manner allows for the greatest predictive power of generated models^37^. Additionally, more rigorous investigation describing the robustness of our models as a function of training and testing sets is found in SI Table S9.

SVRs were performed using various kernels, including Polynomial (degrees 2, 3, 4, and 5), radial base function, and Sigmoid. Using these algorithms, correlation to ΔΔG from predictions by our CSF were improved and now performed similarly to many other literature models.^14^ (Fig. 2C and Table 1) SVR analysis also demonstrated that the BETA_NOV16 score function performed better than the REF2015 score function (Fig. 2C). For GBT, we found that after an exhaustive grid search of tuneable parameters, this technique was the most predictive of all models tested as it produced the highest correlation as well as lowest MAE. Interestingly, our GBT models were invariant to which Rosetta score function was used for simulation as BETA_NOV16 and REF2015 score functions tested identically.

**Table 1.** Machine Learning Models Using the SKEMPI Database.

| Method | R Value | MAE |
| --- | --- | --- |
| FoldX | 0.34 | 1.33 |
| Pred1 | 0.45 | 1.14 |
| BeAtMuSiC | 0.46 | 1.09 |
| Pred2 | 0.54 | 1.07 |
| MLR | 0.31 | 1.18 |
| SVR | 0.53 | 1.09 |
| **SRS2020** | **0.65** | **0.92** |

Alternative methods for predicting $\Delta\Delta G$ using the SKEMPI database. R value is the Pearson correlation coefficient and MAE is mean absolute error in kcal/mol.

To further identify any potential differences between these two models, feature importance analysis was performed. The two models were extremely similar and terms corresponding to phi-psi or rotameric preferences (Fig. 3, Struct. category) were found to be most important. These terms were followed by solvation (Solv.), van der Waals (Atr. and Rep.), the single value of Rosetta total energy (REU), electrostatic (Elec.), and hydrogen bonding terms (H-Bond). Considering that the database is primarily comprised of mutations to alanine, the typical reduction in size and increase in hydrophobicity associated with these changes likely explains the importance of solvation and nonpolar, attractive interactions over hydrogen bonding or electrostatics. A further breakdown of how SRS2020 predicts specific subsets of the SKEMPI2.0 database is found in SI Table S9.

**Fig. 3.**
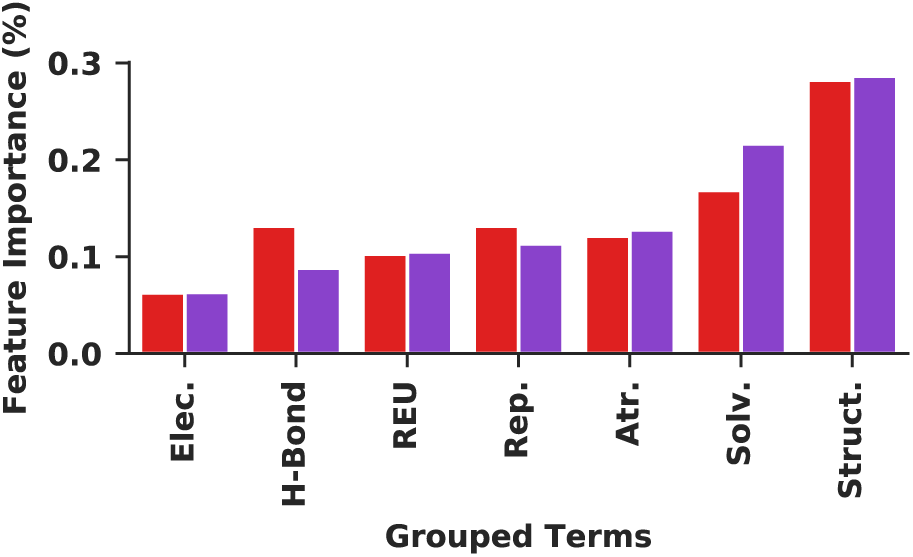
Important features in GBT models derived from REF2015 (red) and BETA_Nov16 (purple). Feature importance (%) determined as described in SI.

After identifying that the GBT-based CSF derived from BETA_NOV16 optimized structures was our best model, we have named it SRS2020 (information on using this code can be found on our github **https://github.com/Sam-Giannakoulias/RML_ddG/** or automated prediction can be performed with our Jupyter web app deposited there. Table 1 compares our results to other machine learning models utilizing a subset of the SKEMPI database to predict ΔΔG. All values in this table represent five-fold cross validation of the exact same subset of single mutations from the SKEMPI database produced by the alternative methods and are thus directly comparable. SRS2020 represents not only the first reported CSF but is the most accurate predictor of ΔΔG in both correlation and in error, highlighting the potential utility of CSF-based approaches. SRS2020 was trained and regressed from the largest protein interaction data set available in the literature and has proven robust to alterations in sampling and scoring, demonstrating the strength of CSF approaches for specific applications. Furthermore, freedom from sampling optimization removes the need to find the perfect simulation and results in an incredibly rapid approach. Prediction of the ΔΔG values upon mutation for a twenty-residue interface can be predicted in less than ten minutes on a single CPU node using this protocol.

We intend to extend this methodology to encompass more complex sampling methods, such as the ensemble-based Backrub approach.^31^ Although this CSF contains no additional energy terms or metrics, one can also easily introduce a variety of bioinformatics terms to further strengthen these models. Additionally, even more complex machine learning methods such as Extreme Gradient Boosted Random Forrest Regressions (XGBoost) or neural networks (NNs)^32^ may be employed to further improve ΔΔG prediction. Finally, we hope to extend the SRS2020 model beyond the prediction of interfacial ΔΔG and use it to design protein-protein interfaces as well as peptides or peptidomimetics targeting such interfaces.

This work was supported by funding from the University of Pennsylvania (UPenn), as well as a grant from the National Science Foundation (NSF CHE-1150351). JJF thanks NSF(DGE-1321851) and the Parkinson’s Disease Foundation (PF-RVSA-SFW-1754) for fellowship support. SRS thanks the UPenn Center for Undergraduate Research & Fellowships for support.

