## Supplementary Information for "Rosetta Custom Score Functions Accurately Predict ΔΔG of Mutations at Protein-Protein Interfaces Using Machine Learning"

**Table of Contents:**

1. **Software…… S2**
2. **Datasets…… S2**
3. **Relax…… S2**
4. **Mutation and Minimization…… S4**
5. **Scoring…… S9**
6. **Multiple Linear Regression…… S12**
7. **Kernel Ridge Regression…… S15**
8. **Support Vector Regression…… S17**
9. **Gradient Boosted Random Forrest Regression…… S18**
10. **Feature Importance…… S20**
11. **Tuning Parameters…… S22**
12. **Cross Validation…… S25**
13. **Comparison to Alternative Methods…… S27**
14. **References…… S28**

### Software

Software necessary for this work includes python3.7, PyRosetta, and the following packages: numpy, scipy, pandas, seaborn, and scikit-learn. Further instruction on how to run these codes can be found at https://github.com/Sam-Giannakoulias/RML_ddG/tree/master/Anaconda where an Anaconda .yml file can be downloaded and used to create a functional virtual environment for this program.

### Datasets

Models generated in this work are derived from a large subset of the SKEMPI2.0 database (https://life.bsc.es/pid/skempi2/).[1] Complexes which contained ambiguous labels of mutational positions were removed leaving a set of 5366 unique mutant complexes. Training and testing datasets for machine learning were curated as to contain no redundant mutations. Further validation of the models using various subsets of the SKEMPI database are shown in the cross validation section. These data sets can be found on our github at https://github.com/Sam-Giannakoulias/RML_ddG/tree/master/Dataset.

### Relax

Wild-type complexes were preprocessed for relax by removing all solvent, ligands, or ions from the structures. Complexes were subsequently renumbered and put into a relax protocol in PyRosetta. The REF2015_CART and BETA_NOV16_CART score functions were used to perform a constrained dualspace relax of the wild-type complexes. Atom pair constraints were set such that $C_\alpha$ atoms found within 8 Å of each other were subjected to a harmonic constraint where the standard deviation of the constraint was 0.5 Å. Additionally, the structures were relaxed with a MoveMap set for both phi/psi and chi angle optimization alongside the minimize bond angles flag.

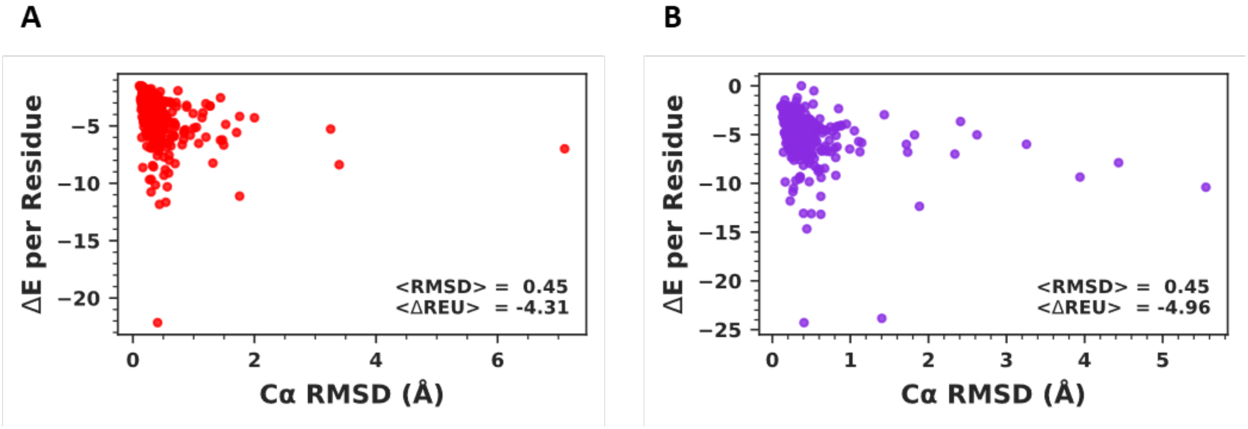


**Fig. S1** Structure and energy differences induced by relax. $C_\alpha$ RMSD (Å) vs normalized Rosetta energy in REU for relaxed structures produced by the (A) REF2015_cart and the (B) BETA_NOV16_cart score functions vs. structures directly from SKEMPI 2.0.

Relaxation of wild-type complexes with both score functions produced little movement of backbone atoms as complexes simulated by both score functions deviated by less than 0.5 Å on average ($C_\alpha$ RMSD, Fig. S1).

$$RMSD = \sqrt{\frac{\sum_i^n (y_i - x_i)^2}{n}}$$

where $y_i$ corresponds to experimental $C_\alpha$ vectors and $x_i$ are altered $C_\alpha$ vectors. Finally, $n$ is the total number of $C_\alpha$ vectors.

Small structural perturbations from the unprocessed SKEMPI complexes were able to substantially decrease the normalized Rosetta energy (Rosetta Energy Units divided by the length of the pose). On average, a given complex displayed a normalized REU decrease of just under five REU, for both score functions.

**Mutation and Minimization**

Mutations of the wild-type complexes were made using the Rosetta Modeling Suite's design functionalities. In simulations where only the mutant residue was packed, resfiles were generated such that the wild-type residues were designed with the NATRO flag to preserve their initial rotameric state. The residue to be mutated was set directly with the PIKAA flag. In simulations where the whole complex was packed, the NATRO flag was replaced with NATAA as to preserve the identity of the residues but optimize their rotameric states. Following packing, minimization was performed using a MinMover set with the following settings: lbfgs_armijo_nonmonotone optimizer, 2000 maximum steps, 0.00001 tolerance threshold, the cartesian flag set to True, and the minimize bond angles flag was set to True. During minimization, a MoveMap was set such that backbone phi and psi angles were not able to be optimized, whereas sidechain chi angles were.

Score files were computed at two stages during the protocol in this work. The first was following relax and the second was after global minimization succeeding mutation (Fig. S2). All score files were recorded using the noncartesian versions of the score functions. Scoring with the REF2015 and BETA_NOV16 score functions as opposed to the REF2015_CART or BETA_NOV16_CART was done to incorporate the effect of the pro_close energy term as well as ignore the cart_bonded energy term (cartesian score functions do not record pro_close and the cart_bonded term is large and highly variable between structures, often misrepresenting the score of the total structure). Mutant scores were subtracted from the wild-type scores generated by identical minimization and packing to produce the inputs for machine learning.

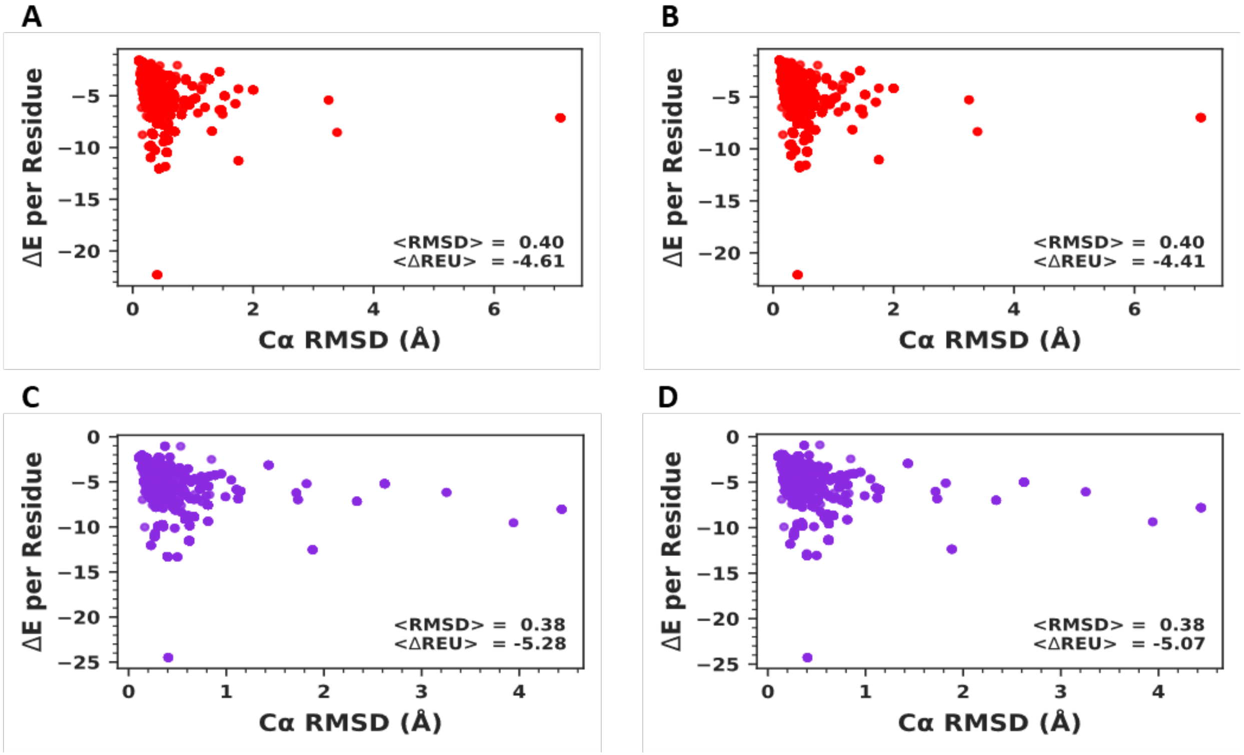


**Fig. S2** Structure and energy differences induced by mutation, packing, and minimization on relaxed complexes. Optimized complexes produced by (A) the REF2015_cart score function and local packing, (B) the REF2015_cart score function and global packing, (C) the BETA_NOV16_cart score function and local packing, or (D) the BETA_NOV16_cart score function and global packing.

After relaxed complexes were mutated and optimized, they again demonstrated small deviation from the unprocessed SKEMPI complexes. In fact, these operations decreased the average Cα RMSD produced by relax for both score functions. Additionally, REU scores for complexes following minimization were decreased even further than during relax in all sampling and scoring schemes. It should be noted that for both score functions, local packing decreased relative REU values further than global packing, as well as that the BETA_NOV16 score function decreased REU values further than the REF2015 score function.

Mutation and optimization of the unprocessed SKEMPI complexes by both REF2015_CART and BETA_NOV16_CART produced a less than 0.01 Å deviation in Cα RMSD. The relative decrease in REU for both score functions at this step was smaller than in the corresponding simulations performed on pre-relaxed structures for both local and global packing prior to minimization. It should be noted that global packing (Fig. S3 B, D) decreased relative REU values further than local packing (Fig. S3 A, C) for unrelaxed complexes, but not for relaxed complexes. This is likely due to the characteristically high score values associated with unrelaxed structures in Rosetta. In all simulations, the BETA_NOV16 score function decreased total REU further than REF2015.

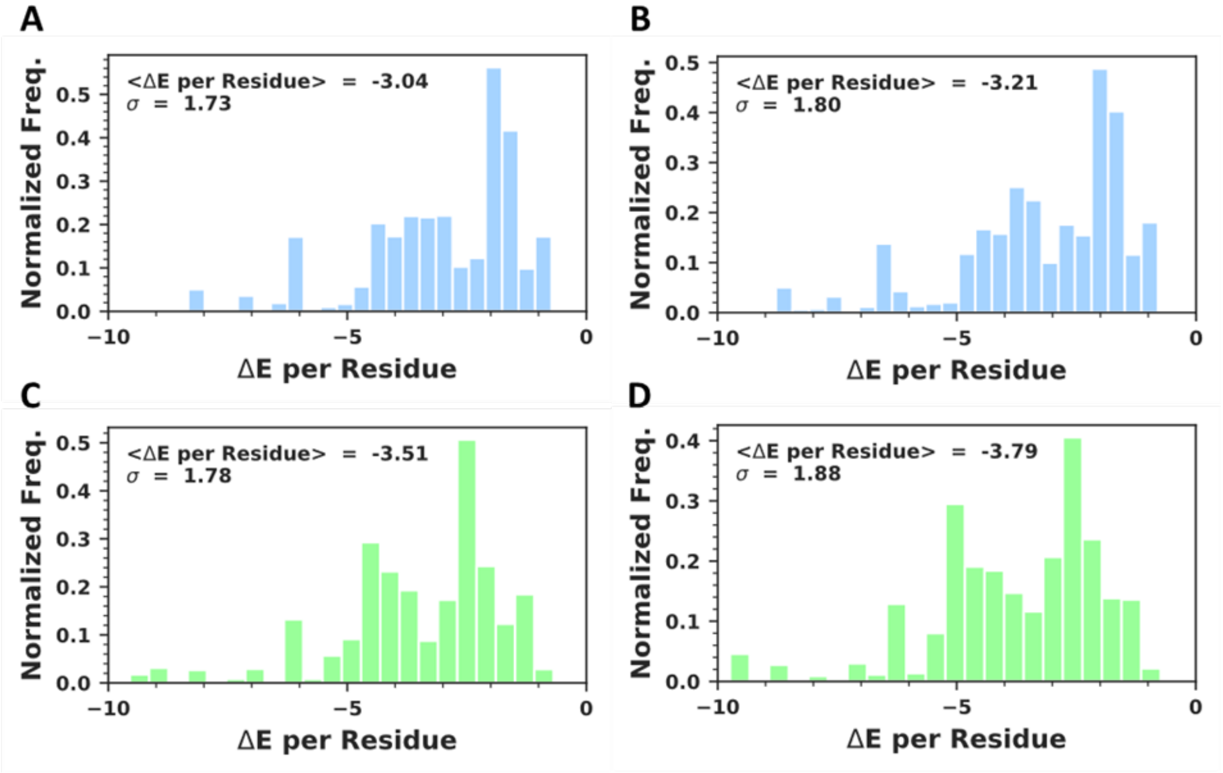


**Fig. S3** Energy difference distribution induced by mutation, packing, and minimization on unrelaxed SKEMPI complexes. Optimized complexes produced by (A) the REF2015_cart score function and local packing, (B) the REF2015_cart score function and global packing, (C) the BETA_NOV16_cart score function and local packing, (D) and the BETA_NOV16_cart score function and global packing.

Differences in sampling schemes produced different distributions in the change in total REU values (Fig. S4). For both the REF2015 and BETA_NOV16 score functions, scores corresponding to local packing generated a narrower energy distribution than scores corresponding to global packing (the standard deviation of global packing simulations was almost two times higher than local packing). There did not appear to be a strong trend in predictive accuracy of ΔΔG and magnitude of change in REU at this step. Global packing simulations with BETA_NOV16 increased relative REU by the smallest margin while local packing simulations with BETA_NOV16 increased relative REU by the largest margin.

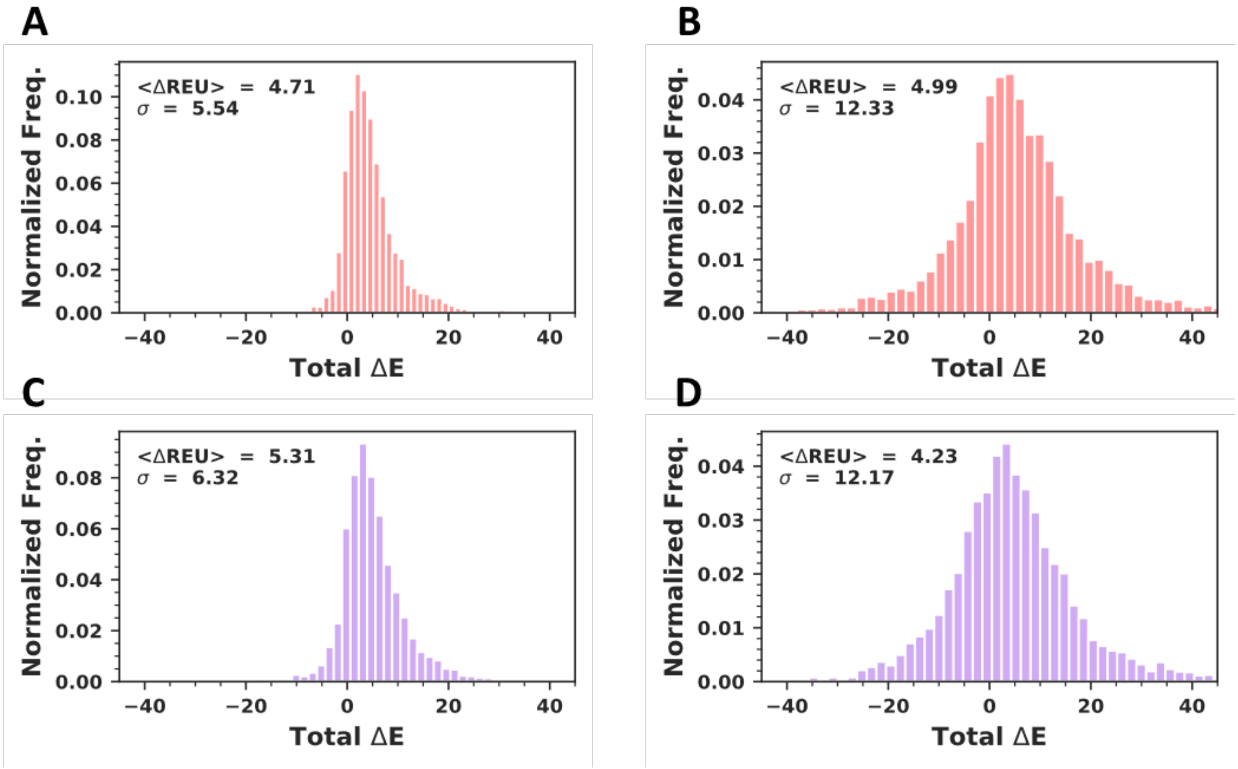


**Fig. S4** Distribution analysis of the change in Rosetta energy as a function of sampling for relaxed complexes**.** Histograms display the distribution of the change in total REU following mutation and minimization compared to the unmutated and minimized relaxed control. (A) shows the REF2015_CART score function and local packing, (B) the REF2015_CART and global packing, (C) the BETA_NOV16_CART score function and local packing, (D) and BETA_NOV16_cart score function and global packing.

Similarly, to the relaxed simulations, distributions of the change in total REU for the unrelaxed complexes displayed the trend that local packing produced a narrower distribution of energies than global packing for both the REF2015 and BETA_NOV16 score functions (Fig. S5). Interestingly, the unrelaxed simulations displayed a different, but equally uncorrelated trend in change in REU and predictive power of ΔΔG. Here, REU values were increased the most by BETA_NOV16 and global simulations and least by REF2015 and global packing simulations.

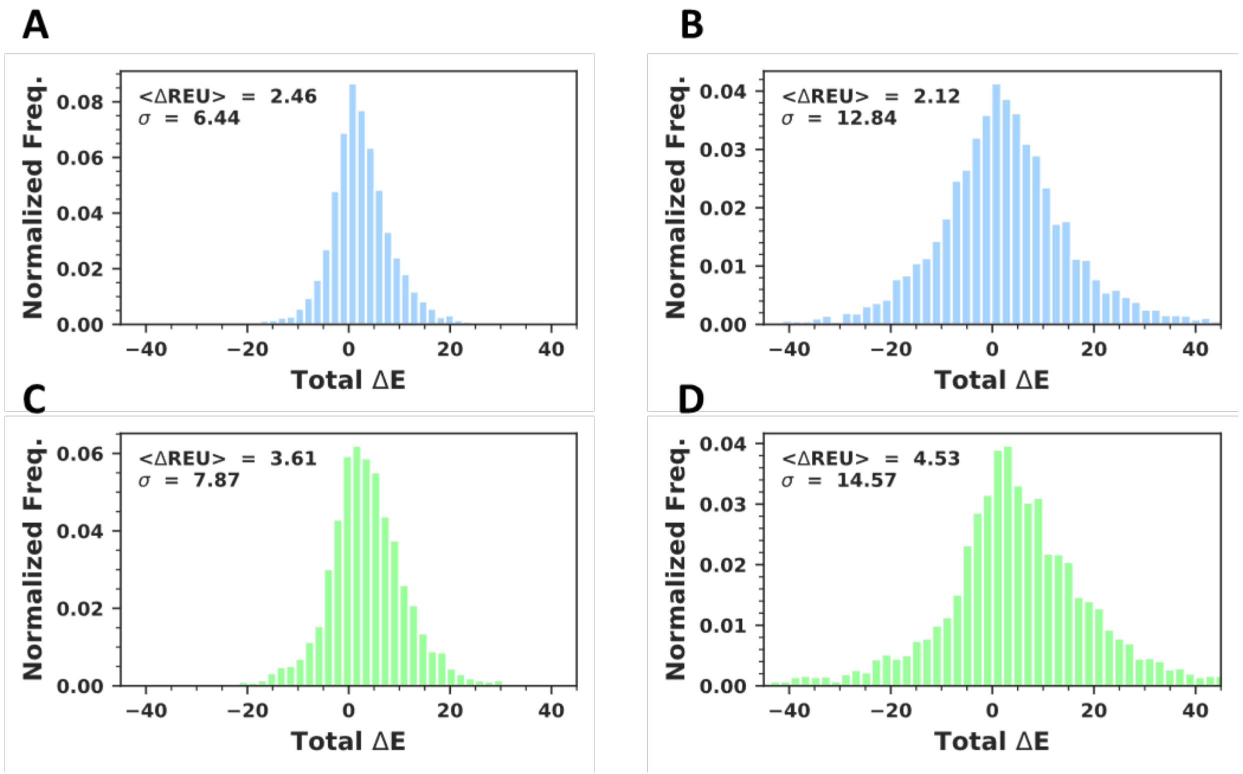


**Fig. S5** Distribution analysis of the change in Rosetta energy as a function of sampling for unrelaxed complexes. Histograms display the distribution of the change in total REU following mutation and minimization compared to the unmutated and minimized unrelaxed control. (A) shows the REF2015_CART score function and local packing, (B) the REF2015_CART and global packing, (C) the BETA_NOV16_CART score function and local packing, (D) and BETA_NOV16_cart score function and global packing.

These data demonstrate that the distributions of energies produced by local packing look much more similar to experimental ΔΔG, the dependent variable, than distributions from global packing. The experimental ΔΔG distribution is shown in Fig. S6 for comparison. This could potentially explain the greater correlation of the local packing simulations as opposed to the global packing simulations.

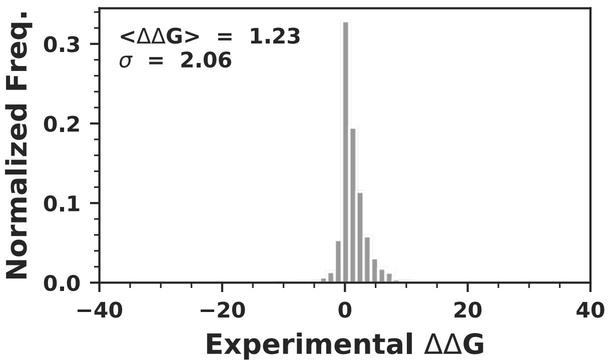


**Fig. S6** Histograms showing the distribution of ΔΔG values in the SKEMPI2.0 database.

### Scoring

To further investigate the differences in predictive capacities of models produced via different sampling schemes, we performed student's, and Wilcoxon t tests on the Rosetta score terms. This analysis is presented in Tables S1 and S2.

**Table S1.** Hypothesis testing of energy terms in the REF2015 local and global packing simulations

| Score Term | Student's t-Test | p-value | Wilcoxon Score | Wilcoxon p-value |
|---|---|---|---|---|
| yhh_planarity | 11.88 | $3.66x10^{-32}$ | 5037353 | $7.25x10^{-28}$ |
| fa_sol | -9.18 | $6.40x10^{-20}$ | 5272429 | $3.82x10^{-21}$ |
| fa_rep | -6.41 | $1.6x10^{-10}$ | 5447204 | $1.15x10^{-14}$ |
| hbond_bb_sc | 6.30 | $3.22x10^{-10}$ | 5889881 | $8.01x10^{-04}$ |
| fa_elec | 6.11 | $1.06x10^{-09}$ | 5694906 | $1.22x10^{-07}$ |
| dslf_fa13 | -5.81 | $6.61x10^{-09}$ | 3379259 | $1.56x10^{-14}$ |
| fa_atr | 4.99 | $6.17x10^{-07}$ | 5703000 | $1.88x10^{-07}$ |
| hbond_sc | -4.14 | $3.45x10^{-05}$ | 5747521 | $2.03x10^{-06}$ |
| pro_close | -3.47 | $5.30x10^{-04}$ | 5401845 | $3.79x10^{-16}$ |

**Table S2.** Hypothesis testing of energy terms in the BETA_NOV16 local and global packing simulations

| Score Term | Student's t-Test | p-value | Wilcoxon Score | Wilcoxon p-value |
|---|---|---|---|---|
| fa_atr | 8.27 | $1.67 \times 10^{-16}$ | 5356365 | $7.09 \times 10^{-18}$ |
| fa_dun_semi | 7.73 | $1.23 \times 10^{-14}$ | 5352622 | $5.14 \times 10^{-18}$ |
| fa_intra_atr | -6.12 | $9.96 \times 10^{-10}$ | 5288207 | $3.04 \times 10^{-20}$ |
| lk_ball_iso | -6.02 | $1.87 \times 10^{-09}$ | 5640469 | $5.73 \times 10^{-09}$ |
| lk_ball | -5.65 | $1.61 \times 10^{-08}$ | 5616268 | $1.34 \times 10^{-09}$ |
| fa_intra_sol | 5.42 | $6.00 \times 10^{-08}$ | 5465204 | $7.53 \times 10^{-14}$ |
| hxl_tors | 4.03 | $5.40 \times 10^{-05}$ | 5961300 | 0.0148 |
| hbond_sr_bb | -3.85 | 0.000119 | 19531 | $6.88 \times 10^{-10}$ |
| dslf_fa13 | -3.05 | 0.002272 | 3069203 | $5.10 \times 10^{-36}$ |
| fa_intra_rep_xover | 2.57 | 0.010038 | 5834168 | 0.000117 |
| fa_dun_rot | 2.46 | 0.013566 | 5809448 | 0.236333 |

These data demonstrate for the REF2015 score function (Table 1) and the BETA_NOV16 score function (Table 2), energy terms that were statistically significantly different from each other between local and global packing simulations via a student's t-test and Wilcoxon test.

In addition to analyzing how Rosetta score terms were different between sampling methods, we also compared how they differed between the score functions themselves. Fig. S7 shows the correlations between the local and global versions of the two score functions for both relaxed and unrelaxed simulations. Fig. S8 shows heatmaps displaying correlations among the simulations and with experimental ΔΔG.

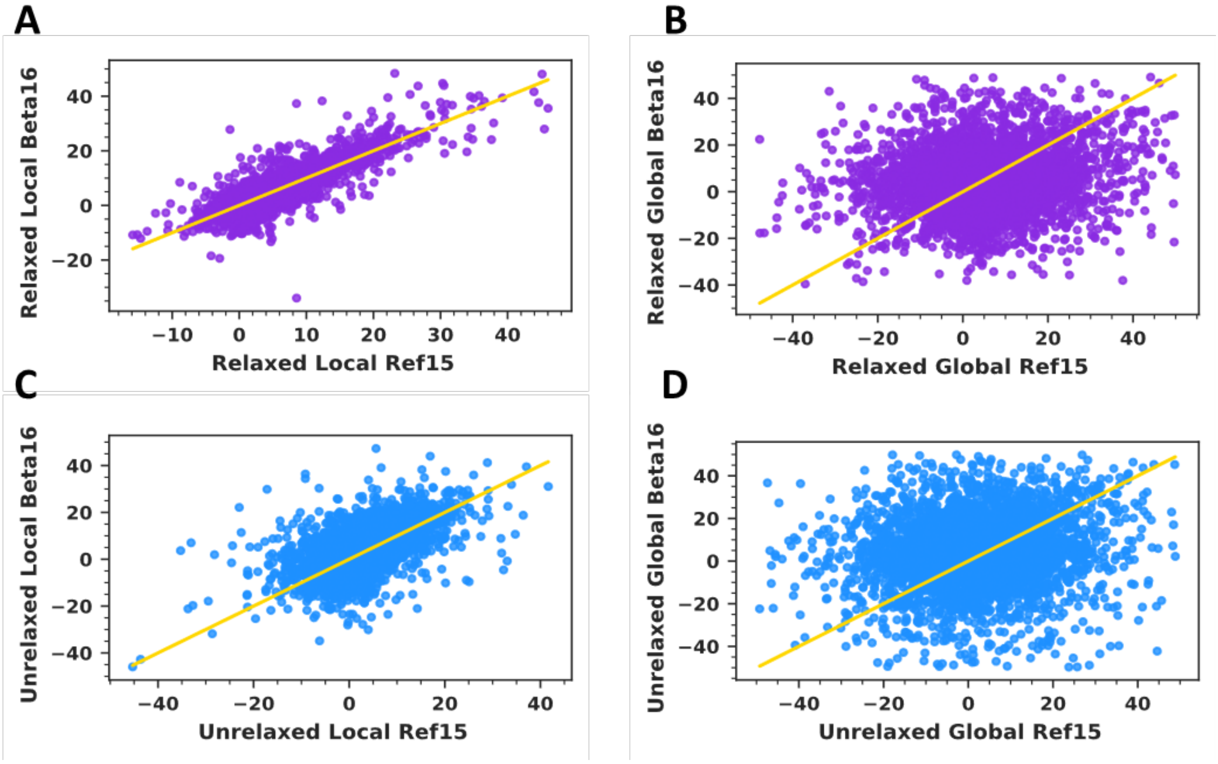


**Figure S7.** Correlation between REF2015 vs. BETA_NOV16 total REU from (A) relaxed local sampling, (B) relaxed global sampling, (C) unrelaxed local sampling, (D) and unrelaxed global sampling (D).

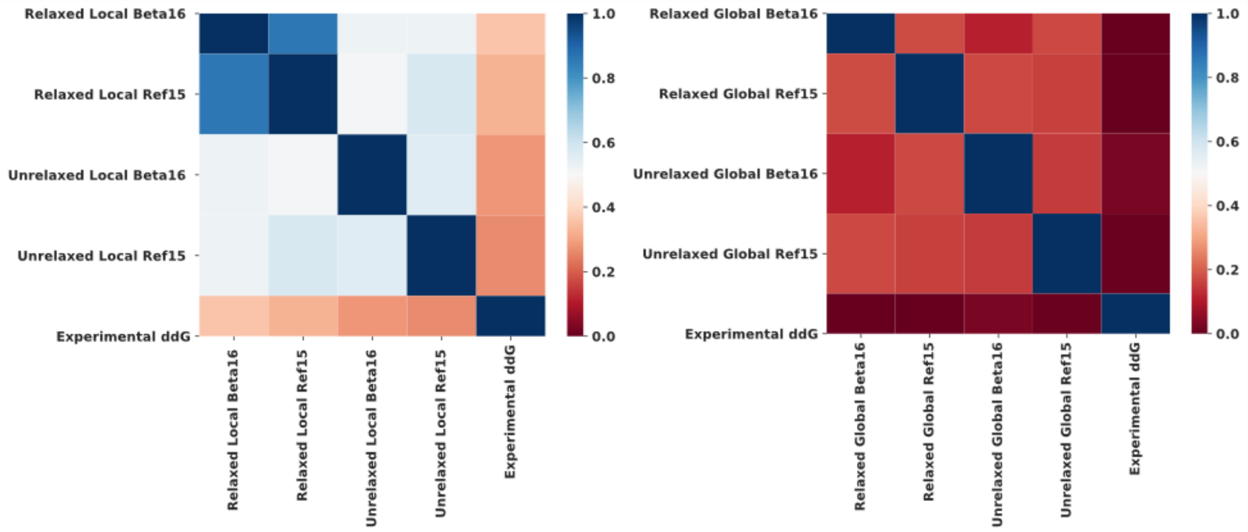


**Fig. S8** Correlation of total REU from all simulations of (A) local packing and (B) global packing with each other.

These data display the correlation between the total REU produced by the REF2015 and BETA_NOV16 score functions as a function of sampling scheme. The difference in correlation between local and global packing simulations was much greater than between simulations of relaxed and unrelaxed complexes. In local packing simulations, which were shown to be more predicative of experimental ΔΔG than global packing simulations, the REF2015 and BETA_NOV16 score functions were found to be quite correlative with each other. This indicates that these score functions are highly similar, but that the BETA_NOV16 score function may be slightly better tuned to energy than REF2015, as it was typically more predicative.

**Multiple Linear Regression (MLR)**

Machine learning algorithms were implemented using scikit-learn. Input features were scaled using a MinMax scaler which can be obtained from sklearn.preprocessing.MinMaxScaler(). Additionally, the input data was processed as to remove outliers or complexes whose simulations did not converge. This process was performed by taking only complexes which fell between -50 and 50 total REU, which corresponded to over 97.5% of the data. Input features were randomly split into 80% training and 20% testing sets for regression using sklearn.model_selection.train_test_split. Metrics such as the coefficient of determination ($R^2$) and mean absolute error (MAE) were computed from scipy.stats.pearsonr and sklearn.metrics.mean_absolute_error respectively.

$$R^2 = \left( \frac{n(\sum_i^n x_i y_i) - (\sum_i^n x_i)(\sum_i^n y_i)}{\sqrt{[n \sum_i^n x^2 - (\sum_i^n x_i)^2][n \sum_i^n y^2 - (\sum_i^n y_i)^2]}} \right)^2$$

$$MAE = \frac{\sum_i^n |x_i - y_i|}{n}$$

where $y_i$ corresponds to experimental values and $x_i$ are predicted values. Finally, $n$ is the total number of data points.

Multiple Linear Regression (MLR) was performed using the sklearn.linear_model.LinearRegression() linear regression module. MLR is a machine learning method which weights many explanatory parameters according to minimization of the least squares criterion.[2] MLR implementation demonstrated that simple reweighting the native Rosetta energy terms can drastically increase the accuracy of prediction of experimental ΔΔG (Fig. S9). Through this method alone, MAE is decreased from over 5 kcal/mol to just over 1 kcal/mol. The general formula for MLR is as follows for a set of k parameters:

$$y = \beta_0 + \beta_1 x_1 + \beta_2 x_2 + \cdots \beta_k x_k.$$

The optimized formulas for our MLR models are shown in Table S3. These weights may be used to create a custom score function (CSF) object in PyRosetta.

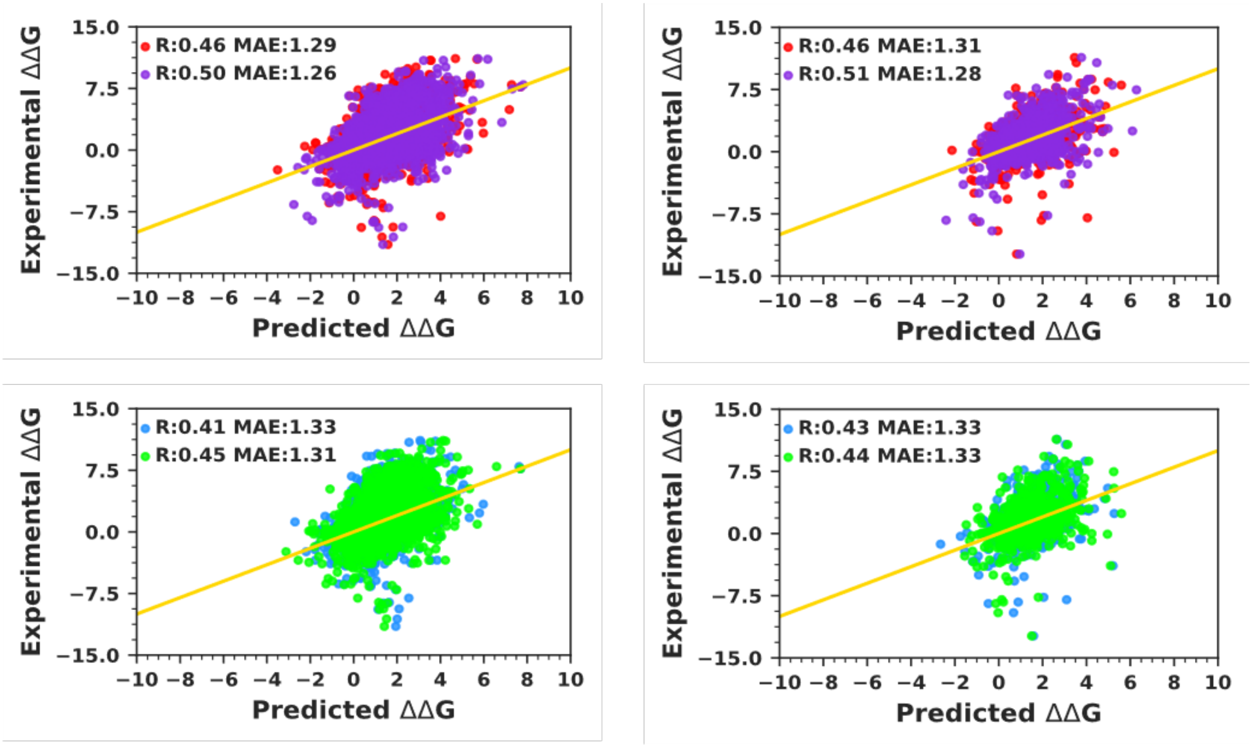


**Fig. S9** Multiple Linear Regression (MLR) predicting ΔΔG. (A and C) Training and (B and D) testing plots of MLR models. Relaxed simulations for REF2015 are in red and relaxed simulations for BETA_NOV16 are in purple. Unrelaxed simulations for REF2015 are in blue and unrelaxed simulations for BETA_NOV16 are in green.

**Table S3.** Weights for CSF based on REF2015 or BETA_NOV16 MLR

| Energy Term | REF2015 Weight | BETA_NOV16 Weight |
|---|---|---|
| fa_atr | 18.27 | 19.144 |
| fa_rep | 4.86 | 4.08 |
| fa_sol | 8.05 | 4.87 |
| fa_intra_atr_xover | N/A | 3.80 |
| fa_intra_rep_xover | 3.21 | 5.00 |
| fa_intra_sol_xover | 2.40 | 2.96 |
| lk_ball | 0.11 | 1.29 |
| lk_ball_iso | N/A | 6.29 |
| lk_ball_bridge | N/A | 0.06 |
| lk_ball_bridge_iso | N/A | -2.18 |
| fa_elec | 9.22 | 6.63 |
| fa_intra_elec | N/A | 1.70 |
| pro_close | 2.58 | 3.38 |
| hbond_sr_bb | -3.98 | -6.00 |
| hbond_lr_bb | 3.92 | 4.70 |
| hbond_bb_sc | -1.48 | -0.54 |
| hbond_sc | 0.55 | 2.25 |
| dslf_fa13 | 3.13 | -0.54 |
| omega | -1.76 | -0.74 |
| fa_dun_dev | 3.92 | 1.14 |
| fa_dun | -2.29 | 3.51 |
| fa_dun_semi | N/A | 2.56 |
| p_aa_pp | 1.23 | -1.66 |
| hxl_tors | N/A | -4.27 |
| ref | -0.61 | -0.24 |
| intercept | -18.24 | -20.97 |

**Kernel Ridge Regression (KRR)**

Kernel Ridge Regression (KRR) was performed using the sklearn.kernel_ridge.KernelRidge() linear regression module. KRR is an effective regression technique which expands upon MLR through the use of the kernel trick and regularization, which penalizes overfitting and creates a more robust model than a MLR. Regularization in KRR differs from regularization in support vector regressions (SVRs) only in the minimization of different loss functions. KRR minimizes squared error loss and utilizes L2 regularization, while SVRs minimize insensitive loss in addition to using L2 regularization.[3] The results of the KRR linear kernel model are shown in Fig. S10. The results of the Laplacian linear kernel model are shown in Fig. S11.

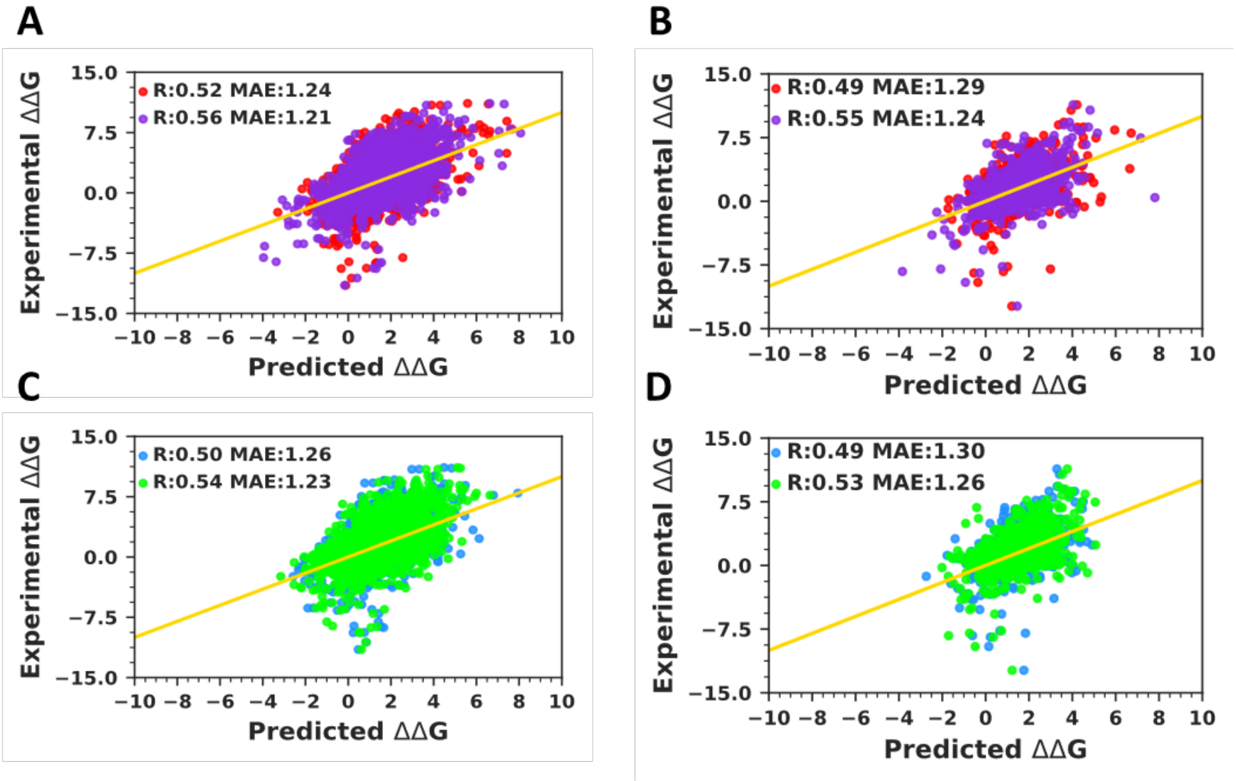


**Fig. S10** Kernel Ridge Regression (KRR) models predicting ΔΔG using a linear kernel. (A and C) Training and (B and D) testing plots. Relaxed simulations for REF2015 are in red and relaxed simulations for BETA_NOV16 are in purple. Unrelaxed simulations for REF2015 are in blue and unrelaxed simulations for BETA_NOV16 are in green.

These data demonstrate that for both the linear and Laplacian kernels, regularization techniques as well as the additional terms corresponding to the energies of the specific mutation site, the 1st contacting shell (8 Å , "local"), and the 2nd contacting shell (16 Å, "distal"), increased testing correlation and decreased testing error modestly.

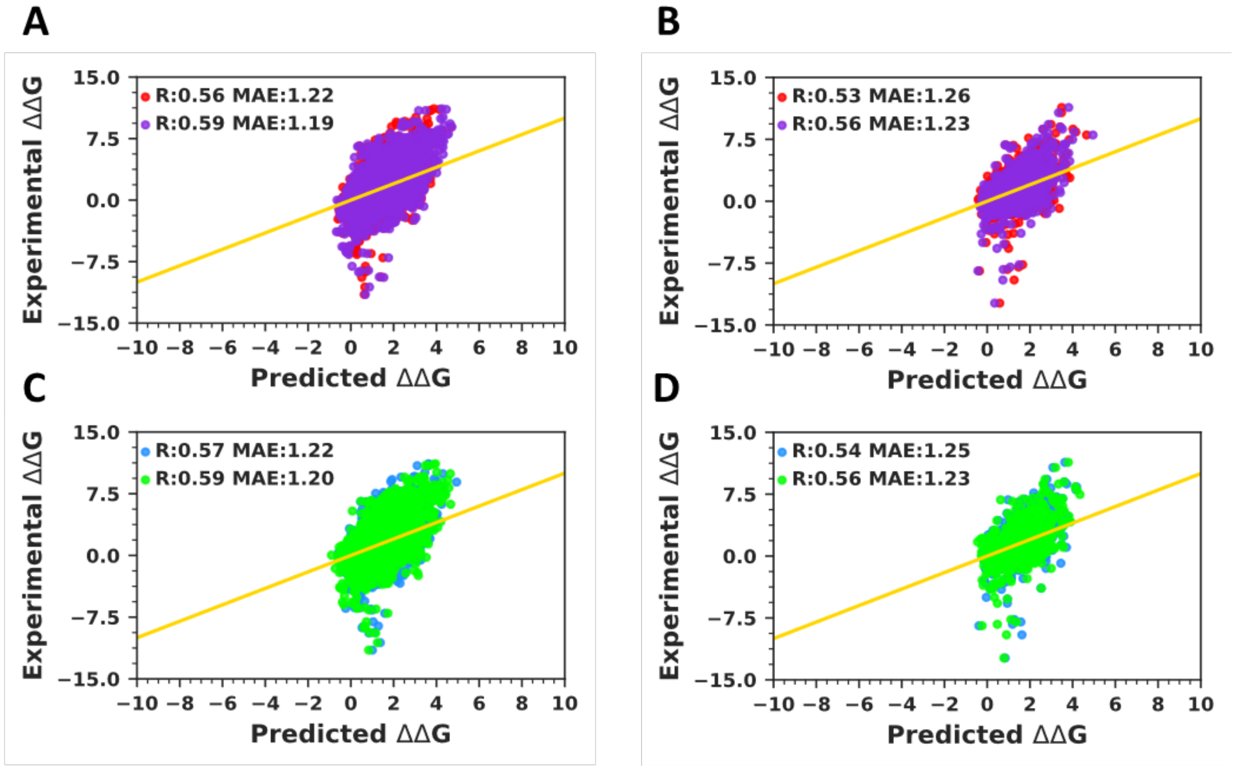


**Figure S11.** KRR models predicting ΔΔG using a Laplacian kernel, (A and C) Training and (B and D) testing plots. Relaxed simulations for REF2015 are in red and relaxed simulations for BETA_NOV16 are in purple. Unrelaxed simulations for REF2015 are in blue and unrelaxed simulations for BETA_NOV16 are in green.

**Support Vector Regressions (SVR)**

Support vector regressions were performed using the sklearn.svm.SVR() module, where the kernels tested include: the radial base function, and polynomial degrees(2-5). SVRs utilize the kernel trick which is used to analyze patterns in the data through an implicit representation of the feature space without an explicit transformation. This allows SVRs to fit data rapidly and accurately. Like KRR, SVRs utilize L2 regularization, but minimize different loss functions. SVRs employ an epsilon insensitive loss function which tolerates error to within a tunable value epsilon. The smaller the value of epsilon, the more accurately the hyperplane fits the data.[4] For small positive values of epsilon, SVRs learn a sparse model and thus predict quickly relative to KRR. The RBF (Fig. S12), and polynomial (Fig. S13) kernels were tested in SVRs to predict ΔΔG. These data demonstrate that both the RBF and polynomial kernels performed strongly, increasing correlation and decreasing MAE more effectively than MLR or KRR.

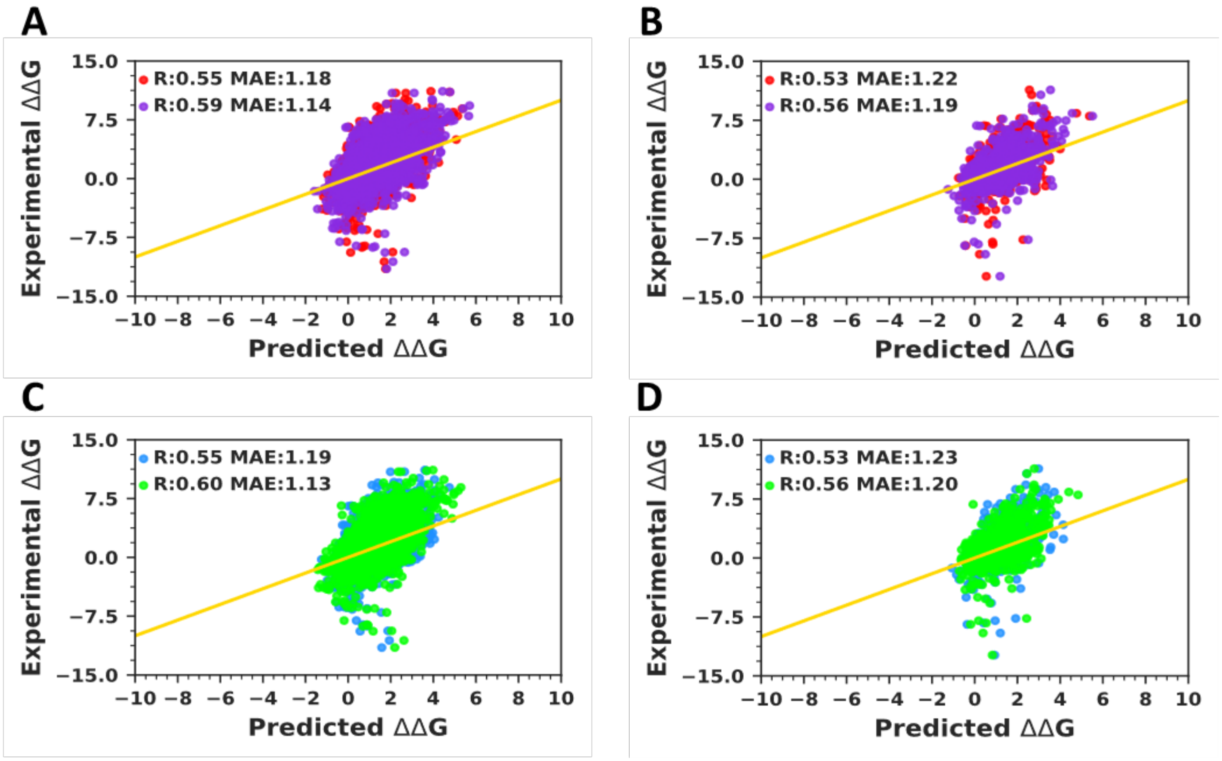


**Fig. S12** SVR models predicting ΔΔG using an RBF kernel, (A and C) Training and (B and D) testing plots. Relaxed simulations for REF2015 are in red and relaxed simulations for BETA_NOV16 are in purple. Unrelaxed simulations for REF2015 are in blue and unrelaxed simulations for BETA_NOV16 are in green.

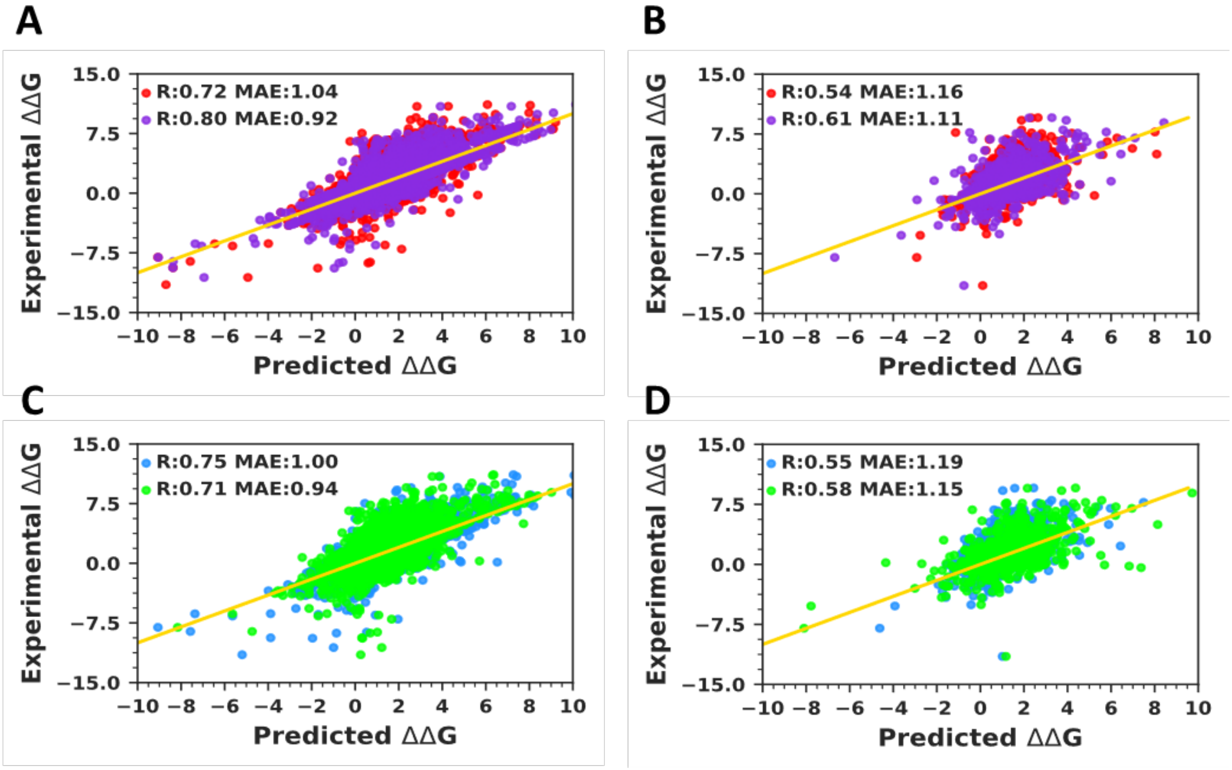


**Fig. S13** SVR models predicting ΔΔG using polynomial kernels, (A and C) Training and (B and D) testing plots. Relaxed simulations for REF2015 are in red and relaxed simulations for BETA_NOV16 are in purple. Unrelaxed simulations for REF2015 are in blue and unrelaxed simulations for BETA_NOV16 are in green.

### Gradient Boosted Random Forrest (GBT)

Gradient Boosted Random Forrest Regression (GBT) was performed using the sklearn.ensemble.GradientBoostingRegressor() module. GBT is an ensemble technique capable of identifying complex patterns within input features, therefore is able to accurately regress relationships. Random forest methods are based on parallelized decision trees, flow chart like structures that use input data to find complex relationships within a dependent variable of interest. Unlike kernel-based methods, the key advantage of tree-based methods is the ability to fit data without an explicit functional form. GBT improve random forest regression through the technique of boosting. Boosting in GBT employs the idea that weak learners (a model which individually is a poor predictor of the dependent variable), can be sequentially used to produce an ensemble model which is a strong learner.[5]

Parameters for GBT in sklearn include loss function, learning rate, number of estimators, subsample, min samples for split, min samples per leaf, min weight fraction per leaf, max depth, max impurity split, random state, max features, and max leaf nodes. To determine the optimal parameters for the models, a cross validated exhaustive grid search of select parameters with sklear.model_selection.GridSearchCV() was utilized. Optimal parameters for SRS_2020 are displayed below in the tuning section.

GBT models were our most predictive models, demonstrating highest correlation and lowest MAE (Fig. S14). These data demonstrate the strength of using GBT for regression using Rosetta energy terms. Relaxed and unrelaxed simulations as well as simulations with both the REF2015 and BETA_NOV16 score functions converged to approximately R 0.75 and MAE of 1.00 respectively.

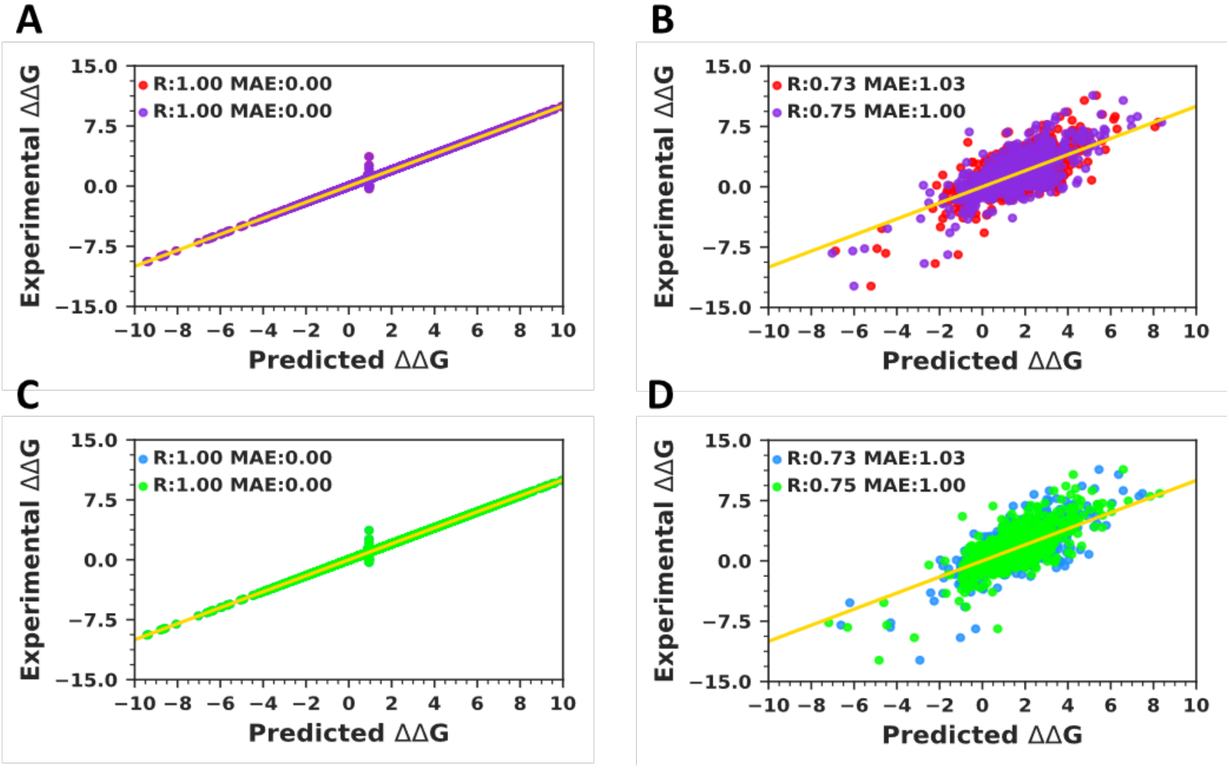


**Fig. S14** GBT models predicting ΔΔG, (A and C) Training and (B and D) testing plots. Relaxed simulations for REF2015 are in red and relaxed simulations for BETA_NOV16 are in purple. Unrelaxed simulations for REF2015 are in blue and unrelaxed simulations for BETA_NOV16 are in green.

### Feature Importance

GBT models were analyzed to asses which features were most relevant in predicting ΔΔG using sklearn. Feature importance metrics can be extracted using the command GradientBoostingRegressor(parameters).feature_importances. The presence of many similar terms such as multiple terms describing solvation or attractive terms found locally, as well as distally, provided an opportunity combine like terms and represent unique categories of features. Table S4 below describes the categorization of these terms. Note, all identical terms only differing by distance were placed in the same category.

**Table S4.** Feature importance classification

| Category | Members |
|---|---|
| Elec (Electrostatic Terms) | fa_elec, fa_intra_elec |
| H-Bond (Hydrogen Bonding Terms) | hbond_sc, hbond_sr_bb, hbond_lr_bb, hbond_bb_sc, |
| REU (Rosetta Total Energy) | total_score |
| Rep (Repulsive Terms) | fa_rep, fa_intra_rep |
| Atr (Attractive Terms) | fa_atr, fa_intra_atr |
| Solv (Solvation Terms) | fa_sol, fa_intra_sol, lk_ball, lk_ball_iso, lk_ball_bridge, lk_ball_bridge_uncpl |
| Struct (Structural Terms) | pro_close, dslf_fa13, rama_prepro, fa_dun, fa_dun_rot_semi, omega, p_aa_pp, ref, hxl_tors |

**Tuning Parameters**

The parameters for all machine learning models were tuned through an exhaustive grid search that is ten-fold cross validated. Final parameters for SRS2020 are displayed here, while all others can be found in csv files uploaded on our github: (**https://github.com/Sam-Giannakoulias/RML_ddG/**).

KRR with a linear kernel takes only the α parameter. The value that the α parameter is tuned to affects the magnitude of the L2 regularization. Regularization seeks to introduce bias into a model with the goal of reducing its overall variance. L2 regularization functions through penalization of the sum of squared weights scaled by the alpha parameter. As α tends towards zero, KRR tends to an ordinary linear regression. KRR can be described by the following equation:

$$\underbrace{\sum_{i=1}^{n}(y_i - \sum_{j=1}^{p} x_{ij}\beta_j)^2}_{\text{Loss Function}} + \underbrace{\alpha \sum_{j=1}^{p} \beta_j^2}_{\text{L2 Regularization}}$$

Here $y_i$ are experimental values, $x_i$ and $\beta_i$ are the features and their corresponding weighting factors respectively, with $\alpha$ finally acting as the scaling factor of the L2 regularization penalty. The optimized parameters are given in Table S5.

**Table S5.** KRR parameters

| Parameter | Min Value | Max Value | Iteration | Final Value |
|---|---|---|---|---|
| α | 0.1 | 10 | 0.1 | 0.1 |

SVR takes two arguments in addition to a kernel. These arguments include a margin of error tolerance parameter (ε) and a regularization parameter (C). SVRs act to minimize error, selecting for hyperplanes (support vector machine) which maximizes the margin (distance between observations), while tolerating error based on epsilon. SVRs can be linear or non-linear depending on the choice of the kernel. Non-linear kernels are used to transform data into higher dimensional space in order to perform the core linear separation. SVRs can be described by the following general form:

$$f(x) = \sum_{i=1}^{n} (\alpha_i - \alpha_i^*) K(x, x_i) + b$$

$$\text{Subject to} \begin{cases} f(x_i) - y_i \leq \varepsilon + \xi_i \\ y_i - f(x_i) \leq \varepsilon + \xi_i^* \\ \xi_i, \xi_i^* \geq 0, i = 1, \dots, n \end{cases}$$

Here, $\alpha_i$ and $\alpha_i^*$ are the Lagrange multipliers afforded through minimization of the regularized cost function subject to ε insensitive loss. The function *K* takes the form of the kernel being tested, b is the bias introduced as in KRR, and the $\xi$ terms are called slack parameters used to assess whether computed hyperplanes fall within the ε threshold.[2, 4] The optimized parameters are given in Table S6.

**Table S6.** SVR parameters

| Parameter | Min Value | Max Value | Iteration | Final Value |
|---|---|---|---|---|
| C | 0.5 | 10 | 0.5 | 10 |
| gamma | Scale | Auto | N/A | scale |
| epsilon | 0.1 | 1.5 | 0.25 | 1 |
| kernel | poly | poly | N/A | poly |
| Degree | 2 | 5 | 1 | 4 |

GBT models utilize many tunable parameters. The loss parameter is defined as the loss function used during regression. Learning rate is a scaling factor which diminishes the contribution of any given tree. The n_estimators parameter describes how many boosting stages should be computed. Min_smaples_leaf is a parameter which describes how many samples are required to be included within the progeny branches at a nodal point in the random forest. The maximum depth directly limits the maximum nodes in the tree. Min samples_split corresponds to the number of samples required for an internal node to be split. Max_features is the maximum number of features which can be considered during each split. Finally, random_state gives the seed produced by a random number generator and is used to incur reproducibility for others. The optimized parameters are given in Table S7.

**Table S7.** GBT parameters

| Parameter | Min Value | Max Value | Iteration | Final Value |
|---|---|---|---|---|
| loss | ls | huber | N/A | ls |
| learning rate | 0.01 | 0.05 | 0.01 | 0.05 |
| n_estimators | 1000 | 1000 | N/A | 1000 |
| min_samples_leaf | 10 | 30 | 5 | 10 |
| max_depth | 5 | 25 | 5 | 10 |
| min_smaples_split | 10 | 50 | 10 | 50 |
| max_features | sqrt or log2 | 0.3 | N/A | sqrt |
| random state | 23 | 23 | N/A | 23 |

These data show the results of cross validation during exhaustive parameter grid search of models generated using energy terms from the BETA_NOV16 score function and local packing. Mean train and test score correspond to the mean Pearson correlation identified via the ten-fold cross validation. These data additionally support that the GBT was our strongest model and that nonlinear techniques were better able to model changes in ΔΔG than linear techniques (Table S8).

**Table S8.** Model evaluation

| Model | Mean train score (CV 10 R) | Mean test score (CV 10 R) |
|---|---|---|
| KRR | 0.318 | 0.276 |
| SVR | 0.375 | 0.322 |
| GBT | 0.998 | 0.532 |

**Cross Validation**

In Table S9 below, we present an analysis of how SRS2020 as well as the polynomial degree three SVR CSF perform on different subsets of the database that they were trained on using five and ten-fold cross validation. We demonstrate that in spite of the over-representation of alanine mutations in the dataset, our models performed best on the subset of data excluding mutations to alanine, although this data was very similar to our full data set (0.757 compared to 0.727 respectively). Additionally, SRS2020 tended to perform better on samples corresponding to multiple mutations more effectively than single mutations. This was also observed by the pred2 score function produced by Li *et. al*.[6] Finally, to rigorously test the robustness of SRS2020, we created a subset of completely unique residue sites by leaving at most one mutation for any given residue. If there were more than one mutation, the one chosen to remain was done so randomly. This test does not allow for any redundancies in mutation or in residue site to be mutated. This class called nonredundant sites performed almost identically to our full set model (0.712 compared to 0.727), indicating that SRS2020 is a robust model.

**Table S9.** CSF cross validation.

| Model | Subset | R | MAE | CV Num. |
|---|---|---|---|---|
| SVR BETA | Single Mutations | 0.457066141 | 1.03 | 10 |
| SVR BETA | Single Mutations | 0.450602035 | 1.04 | 5 |
| SRS2020 | Single Mutations | 0.583701528 | 0.93 | 10 |
| SRS2020 | Single Mutations | 0.590185202 | 0.93 | 5 |
| SVR BETA | Multiple Mutations | 0.667945297 | 1.51 | 10 |
| SVR BETA | Multiple Mutations | 0.648026151 | 1.58 | 5 |
| SRS2020 | Multiple Mutations | 0.779840086 | 1.26 | 10 |
| SRS2020 | Multiple Mutations | 0.774010895 | 1.28 | 5 |
| SVR BETA | Alanine Only | 0.580558951 | 0.97 | 10 |
| SVR BETA | Alanine Only | 0.578787913 | 0.97 | 5 |
| SRS2020 | Alanine Only | 0.65315039 | 0.9 | 10 |
| SRS2020 | Alanine Only | 0.640055599 | 0.92 | 5 |
| SVR BETA | Alanine Excluded | 0.625189157 | 1.32 | 10 |
| SVR BETA | Alanine Excluded | 0.641047199 | 1.32 | 5 |
| SRS2020 | Alanine Excluded | 0.757523473 | 1.11 | 10 |
| SRS2020 | Alanine Excluded | 0.749314051 | 1.13 | 5 |
| SVR BETA | Nonredudant Sites | 0.588418261 | 1.19 | 10 |
| SVR BETA | Nonredudant Sites | 0.588984757 | 1.19 | 5 |
| SRS2020 | Nonredudant Sites | 0.71226717 | 1.03 | 10 |
| SRS2020 | Nonredudant Sites | 0.695329568 | 1.04 | 5 |
| SVR BETA | Full Set | 0.601991903 | 1.15 | 10 |
| SVR BETA | Full Set | 0.622135583 | 1.16 | 5 |
| SRS2020 | Full Set | 0.727508770 | 1.02 | 10 |
| SRS2020 | Full Set | 0.719102394 | 1.01 | 5 |

**Comparison to Alternative Methods**

SRS2020 was benchmarked against alterative scoring functions for predicting ΔΔG values from mutations found in a subset of the SKEMPI database. Data sets and predictions of the data sets for the alternative methods was obtained from Li *et. al.*[6] FoldX, BeAtMuSiC, Pred1, and Pred2 were compared through five-fold cross validation of the same subset of SKEMPI data. When we attempted to compare SRS2020 to these methods, we found that a small number (104, < 6%) fell outside of the energy range between -50 and 50 REU. We trimmed the data so that the subset was comparable to all score functions and performed the cross validation. In this direct comparison of score functions SRS2020 was found to produce the highest Pearson correlation and lowest MAE.

**Table S10.** Comparison of Score Function Methods on Single Mutant Subset of SKEMPI utilizing CV 5

| Method | R Value | MAE (kcal/mol) |
|---|---|---|
| FoldX | 0.34 | 1.33 |
| Pred1 | 0.45 | 1.14 |
| BeAtMuSiC | 0.46 | 1.09 |
| Pred2 | 0.54 | 1.07 |
| SRS2020 | 0.65 | 0.92 |
